# A cross-sectional study of the commercial plant-based landscape across the US, UK and Canada

**DOI:** 10.1101/2022.04.08.487708

**Authors:** Nicola Guess, Kevin Klatt, Dorothy Wei, Eric Williamson, Ilayda Ulgenalp, Ornella Trinidade, Eslem Kusaslan, Azize Yilidrim, Charlotte Gowers, Robert Guard, Chris Mills

## Abstract

As plant-based foods comprise an ever-increasing proportion of the diet, understanding the nutritional composition of these products is critical. In this study we assess the nutritional content of all commercial plant-based products across multiple sectors (supermarkets, fast food & sit down restaurants, food delivery companies and manufacturers) in the US, UK and Canada. We identified 3488 unique products. Across all sectors, 45% of main meals had >15g protein, 60% had <10%kcal from saturated fat; 29% had >10g fibre per meal; 86% had <1000mg sodium. At restaurants, meat-based main meals were significantly higher in protein and sodium compared to vegetarian and vegan meals. The meat-based options were also significantly higher in saturated fat than the vegan but not vegetarian options. We conclude that plant-based items tend to be lower in saturated fat and sodium than their meat-based counterparts but improvements are needed to optimise their nutritional composition.

## Introduction

There has been a rapid increase in the consumption of plant-based alternatives over the past decade. The worldwide plant-based meat market was US$ 5.6 Billion in 2020 with an expected growth rate of 15% between 2020-2027 [1]. Sales of plant-based foods in the UK increased by 40% between 2014 and 2019 [2] and in 2018, 16% of new food launches in the UK were vegan [3]. Nearly one in four Americans (23%) report eating less meat in the past year than they had previously according to a 2020 Gallup poll [4].

Animal-based products are large determinants of dietary protein and several micronutrient, in addition to containing nutrients of public health concern including saturated fat and sodium in the diet [5, 6]. Thus, the impact of replacing animal-based products with plant-based alternatives has potential implications for both nutrient adequacy and chronic disease risk factors.

Sufficient dietary protein intake is essential for growth, development and well-being, including the maintenance of a healthy immune system [8]. National and international bodies recommend a daily protein intake of 0.8g protein per kilogram (kg) of body weight [8]. Animal-based products are by far the largest contributor to total protein intake in Western populations [9, 10]. While protein requirements can be met with a completely plant-based (i.e. vegan) diet [11, 12], protein insufficiency is more likely with plant-based diets [12].

Animal products also contribute at least 75% of daily saturated fat intake [13]. A switch from animal to plant-based proteins potentially offers a route to lowering population saturated fat intakes [14]. However, some plant-based meat alternatives are formulated with coconut or palm oil and the potential contribution of plant-based alternatives and meals to saturated fat intakes is unknown. While animal products lack dietary fibers, the composition of plant-based alternatives can vary depending on the degree of refining and texturization of plant proteins, and thus, replacement options may contribute minimal, good or excellent sources of dietary fibers [14]. While plant foods are typically naturally low in sodium, processed plant-based meals may be high in sodium [7].

The net impact of replacing animal-based options with plant-based alternatives will ultimately depend on the specific replacement that occurs, but it remains unclear how the existing marketplace of plant-based products influences these key nutrients of concern, on average. To date, work to identify the nutritional composition of plant-based alternatives has tended to focus on narrow sectors [15], a limited range of products [6, 16, 17] or focussed on narrow geographical areas [7, 15].

While supermarket-based purchases contribute the largest proportion to average intake per capita, outside-of-home food consumption is also a significant contributor [18-22]. In the UK, 20-25% of adults eat meals prepared out-of-home weekly [18]. In the US in 2018 44% of all food spending was away from home [21]. 30% of USA adults consume 5 or more meals away from home per week [19]. Other analyses across five countries suggest 78% had purchased at least one meal prepared away-from-home in the past 7 days [20].

Therefore, any analysis of the nutritional quality of plant-based products must take into account the most common providers of food. To that end, we sought to characterize the existing commercial landscape of products marketed as plant-based alternatives and document their available nutrient composition data across countries (USA, UK, Canada), procurement sectors, and meal categories. These efforts provide a snapshot of the nutritional landscape of the existing plant-based alternative market and serves to both help further refine clinical and public health guidance and inform manufacturers when formulating new products.

## Methods

### Search and Retrieval Strategy

To assess the current plant-based alternative landscape, we utilized a tiered approach, searching across pre-defined procurement categories in the United States, the United Kingdom and Canada from April 2020 through December 2020; additional product updates occurred throughout quality control inspection of the data through April 2021. Procurement categories included 1) supermarkets, 2) fast food restaurants, 3) sit-down restaurants, 4) meal delivery and 5) manufacturers. In addition, we had intended to collect and analyse data from independent restaurants, hotel and aeroplane menus.

However, due to the COVID19 pandemic, the menus (where available on a website) were limited in product number and diversity for these procurement categories. Furthermore, online nutritional information was not available for any of these products, and our protocol was to contact the company to request this information. Due to widespread closures and staff furlough it was not possible to collect nutritional information for any of the products in these categories. Therefore, we opted not to include these three sectors. Instead, in order to understand more about the nutritional content of plant-based offerings for people seeking such products we decided to assess the nutritional composition of plant-based meal delivery companies.

Given the distinct procurement markets surveyed, various strategies for identifying products marketed as plant-based were tailored to each procurement category.

#### Supermarkets

We identified the top 10 supermarkets based on the most recent market share estimates per country. We searched all supermarket websites using the following search terms: “vegan”, “vegetarian”, “plant-based” “plant-predominant”, “meatless”, “meat free”, “meat replacement”, “meat alternative”, “dairy alternative”.

#### Fast-food and Sit-down Restaurants

We identified the top ten fast food and sit-down restaurant chains per market share in each country. The menus were searched based on whether any item was advertised as vegan, vegetarian or plant-based, whether with words or an icon.

#### Meal Delivery Companies

To represent the likely behaviour of an individual seeking to purchase a plant-based meal for delivery, we input the search terms “plant-based vegan vegetarian delivery” into the Google search engine, using the largest city in each country as the user location. We then extracted the data from the top ten delivery chains for each country in order of appearance. This approach also avoided the duplication of data collection from many of the fast-food and sit-down restaurants which either offered delivery already or commenced offering delivery during the COVID19 pandemic.

#### Manufacturer

In order to capture the nutritional data from the most exhaustive list of plant-based foods possible, whenever we found a branded plant-based item within supermarkets, we then went to the brand’s website and extracted all products that were currently available.

As different fast-food, sit-down and delivery companies used different ways to highlight their plant-based offerings, we used the following strategy: For chains which specifically highlighted their plant-based products using an icon or key on the menu, we included all products highlighted, again coding whether the product was vegetarian or vegan. Across all categories, we sought out products advertised as plant-based alternatives; thus, we intentionally sought out not to capture all products that could be technically consumed on a plant-based diet (e.g. bagged/canned beans at supermarkets; cooked potato side dishes at restaurants).

To assess the nutrient content of the current plant-based product landscape, we relied on nutrition contents provided on the nutrition facts panel, labels and/or menus provided by the individual commercial entity. To be included in the final product database, kilocalorie and macronutrients per unit serving size had to be provided. We also extracted saturated fat, fibre, sugar and sodium information but these nutrients were not available for all products. Where possible, total serving size, suggested serving size, and serving size per 100g were retrieved and/or calculated. Due to country-level differences in the labeling of sodium vs salt, salt values were converted to sodium (salt (g) x 1000 *2.5).

### Data Processing and Analysis

There was a large variety of products available across the various procurement categories. Therefore, to further subcategorize products and identify nutrient composition trends specific to individual products and their intended uses, we categorised the products into separate meal categories: 1) Product intended to be a whole meal (Whole Meal); 2) Main Protein Source; 3) Starter; 4) Side and/or sharing plate; 5) Snacks; 6) Dessert, 7) Sauces/Condiments; 8) Dairy Alternatives. Categories were not mutually exclusive; thus, some products were included in multiple categories. Categorizations prioritized the marketing strategy used for the product (i.e. sold in the menu section for mains, proteins, sides, etc), with additional categorizations based on readily accepted cultural food norms. To ensure representative categorizations, products were first coded into these 8 meal categories by 1 reviewer (NG) and a subset of 200 items (identified by random number generator) were coded by a 2nd reviewer (KCK). Concordance between review categories were >95% on first pass and discordant categorizations were subsequently discussed and harmonized between reviewers. Discrepancy themes emerged, resulting primarily from country-specific differences in perceptions of foods as meals vs side dishes and second pass of the product database was undertaken to update categorizations for similar products. Specific criteria for assigning a product into a meal category are detailed below:

#### Whole meal

For all categories including supermarkets, fast food chains, sit down chains and meal delivery, items were coded as a main meal if they were described as “meal”, “main(s), “entree”, “breakfast”, “lunch” or “dinner”. Where there was no product description, a subjective decision was made as to whether that item might be considered a whole meal. As described above, discrepancies between items were resolved via discussion. An example of a discrepant item in this category was macaroni and cheese which was ultimately coded as both a whole meal and a side dish.

#### Main protein

This was an umbrella category that aimed to capture all products which were marketed as the main protein component of a meal, whether by the manufacturer, supermarket, restaurant or delivery company or could reasonably be considered the main protein by virtue of being a “meat alternative”. Products coded as meat alternatives were predominantly comprised of plant-based alternatives that specifically marketed themselves with a word or morpheme reflective of an animal protein; for example, products utilized both ‘beef’ and ‘beeph’ or ‘chicken’ and ‘chik’n’ were included in this category. A list of all terms used to identify meat-alternatives/main proteins are included in the Supplemental Data file.

For the other products included in this category we assessed whether a product was being marketed as the protein component of the meal by how it was presented or described within the packaging, menus or website. For example, the following were coded as main protein sources: products advertised on menus in entree categories alongside main meat products; products which were shown on the packaging as being served with a starch and/or vegetables/salad; products for which the “serve with” recommendations included a starch and/or vegetables/salad; tofu- and tempeh-based items that were not whole meals.

#### Starter

Starters were coded as such if they were labelled as “appetizers” or “starters” on the menu or website. Where there was no description, subjective judgement was used based on usual intake of these items. For example, soups and salads (except where explicitly noted as “mains”) were coded as starters irrespective of where they appeared in the menu.

#### Side/sharer

This was a combined category to reflect items which could be served as sides or shared plates, or a combination of two.

#### Dairy alternatives

Products were coded as dairy alternatives if they specifically marketed themselves with a word or morpheme reflective of milk, yogurt or cheese. Morphemes for this category can be found in the Supplemental data file. Products were also coded into this category if they were specifically advertised as a dairy/dairy-free alternative, and/or if they were specifically included in the dairy/milk section of a menu and/or supermarket website.

#### Snack

Products were coded as snacks if they contained keywords related to the snack category, including bars, balls, rolls, bites, jerky and chips and other products categorized as ‘crunchy’ or ‘crispy’ (e.g. roasted chickpeas, veggie straws, beetroot crisps). Products that were single-serving, pre-packaged items were commonly coded as snacks unless the met the ‘whole meal’ or ‘main protein’. Products were coded as snacks if they were specifically advertised as snacks, and/or were included in the snack/cupboard section of websites.

#### Dessert

Dessert products were coded as such if they contained keywords related to the dessert category, including ice cream, cake, brownies, caramels, chocolate bars, cupcakes, flavored biscuits, cookies or cookie dough, or pies. Products were also coded as desserts if they were specifically advertised as a dessert product and/or were included in the dessert section of menus and/or supermarket websites.

#### Sauces, and Condiments

Products were coded as sauces, condiments and dips if they contained keywords, including mayo/mayonnaise, dips, sauces, dressing, and pesto. Products were also coded into this category if they were specifically advertised as such, and/or were included in the sauces, condiments or dips section of a menu and/or supermarket website.

#### Quality control

Given the large number of products retrieved in the database, we employed several quality control steps to ensure accuracy of our search and nutrient information. Retrieved products that were advertised as plant-based but are not plant-based specialty products and/or alternatives to animal-based products were manually removed from the database (e.g. mashed potatoes; baby carrots). Foods were first searched, on a per meal category basis, to ensure the macronutrient values were multiplied by their general calorie content (4:4:9 for carbohydrate:protein:fat) and compared to the retrieved calorie information; calculated calories that were <90% or >110% of the label calories were re-checked against their online nutrient facts information and input errors corrected. Nutrient values (kcals, saturated fat, fibre and sodium) were additionally sorted by quintiles, and the highest and lowest quintiles were manually checked for implausible values for each. As part of quality control, we additionally filtered out duplicate items in the database; only duplicates within a country were removed to allow for capturing of country-specific differences in nutrient content due to different formulations, as well as representative portrayal of country-level statistics. Several strategies were taken to identify duplicates. First, MATCH functions were coded in Excel to identify products with similar names; duplicate products were removed manually upon inspection. This was iteratively performed until the MATCH function yielded the closest product that was not a duplicate. MATCH functions were then applied to numeric values, including the percentage of calories coming from fat * the percentage of calories coming from protein, as well as the percentage of calories coming from carbohydrate, and duplicate products were identified and removed. Throughout this process, we noted that, unsurprisingly, branded items were the most common sources of duplicates within countries (e.g. the same brand/flavor of almond milk was identified at multiple supermarkets). Thus, our last duplicate removal step included manual searches of the database, filtered by product brand and procurement categories of major brands/chains, and removed additional duplicates. Products with missing nutrient values were re-queried as the last quality control step prior to analyses and updated if nutrient content information was available.

#### Quantification

As our aims were explicitly descriptive in nature and absent null hypotheses, our primarily analysis presents descriptive summary statistics. ANOVAs were used for the secondary analysis to compare nutritional content of meat, vegetarian and vegan meals. All descriptive summary statistics and graphing were generated in the R Programming Language (R 4.1.0) using the *dplyr* and *ggplot2* packages, respectively.

## Results

Upon completion of search, retrieval and data entry, the plant-based product database contained 4472 products. Following quality control, including removal of non-alternative products that were retrieved (e.g. potatoes; frozen vegetables), removal of within-country duplicates, and removal of products without complete macronutrient data, 3488 products were included in the final analysis (Table 1).

**Table 1:** Number of vegan and vegetarian products found per meal and procurement category, per country.

|  | Vegetarian |  |  | Vegan |  |  |
| --- | --- | --- | --- | --- | --- | --- |
|  | USA | UK | Canada | USA | UK | Canada |
| **Procurement Category** |  |  |  |  |  |  |
| Supermarket | 57 | 160 | 25 | 690 | 1096 | 336 |
| Fast Food | 39 | 2 | 18 | 17 | 10 | 4 |
| Sit-down | 72 | 44 | 42 | 3 | 38 | 20 |
| Manufacturer | 7 | 14 | 5 | 78 | 186 | 161 |
| Delivery | 36 | 21 | 0 | 175 | 119 | 13 |
| **Meal Category** |  |  |  |  |  |  |
| Whole Meal | 134 | 80 | 51 | 268 | 355 | 74 |
| Main Protein Source | 38 | 108 | 17 | 227 | 544 | 203 |
| Meat Alternative | 32 | 59 | 13 | 164 | 360 | 143 |
| Starter | 26 | 14 | 14 | 13 | 31 | 15 |
| Sides and Sharers | 33 | 60 | 18 | 71 | 216 | 45 |
| Dairy Alternatives | 5 | 1 | 2 | 322 | 211 | 164 |
| Snacks | 2 | 7 |  | 52 | 100 | 36 |
| Deserts |  | 11 | 5 | 61 | 165 | 38 |
| Sauces & Condiments | 5 | 7 | 1 | 31 | 100 | 16 |

Supermarkets represented the vast majority of the vegan items found in all countries (USA: 692/967 =72%; UK: 1112/1465 = 76%; Canada: 340/540 = 63%) (Table 1). Availability of vegan items in the top ten fast food and sit-down chains was limited. In Canada there were only 4 vegan items across the 10 fast food restaurants; 10 in the UK and 18 in the US were identified. For sit down restaurants, there were only 3 vegan items available in the US, 20 in Canada and 38 in the UK.

We found a total of 974 items in the main protein source category, 667 items in the meat alternative category, 697 main meals and 697 dairy alternatives.

### Nutrition

Across meal categories, we noted striking similarity between the USA, UK and Canada in nutrition facts, suggestive of generally similar provisions on average across countries; **Figure 1** illustrates this similarity, depicting the broadly similar average macronutrient composition observed for meal categories across countries.

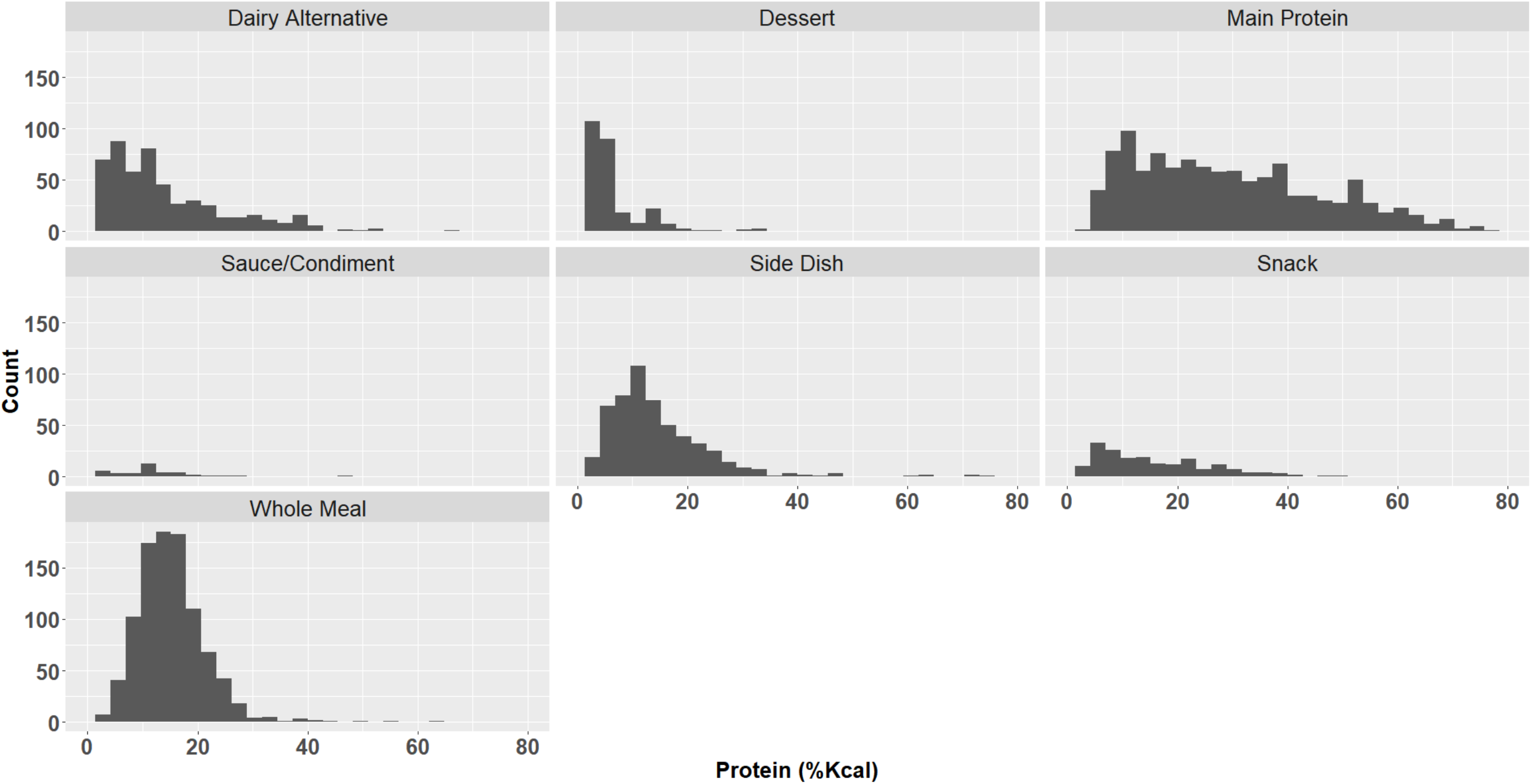

#### Energy

The energy content was available for all 3488 products. The mean energy content of main meals varied from 291.2 ± 135.7kcal (manufacturer) to 587.9 ± 274.7kcal (sit-down restaurants), while the energy content of the main protein ranged from 181.4 ± 96.2kcal (supermarket) to 549.9 ± 63.6kcal (fast food) (Table 2). The energy content of sides/sharers ranged from 201.5 ± 118.8kcal (supermarket) to 534.5 ± 408.5kcal (sit-down restaurant).

**Table 2:**
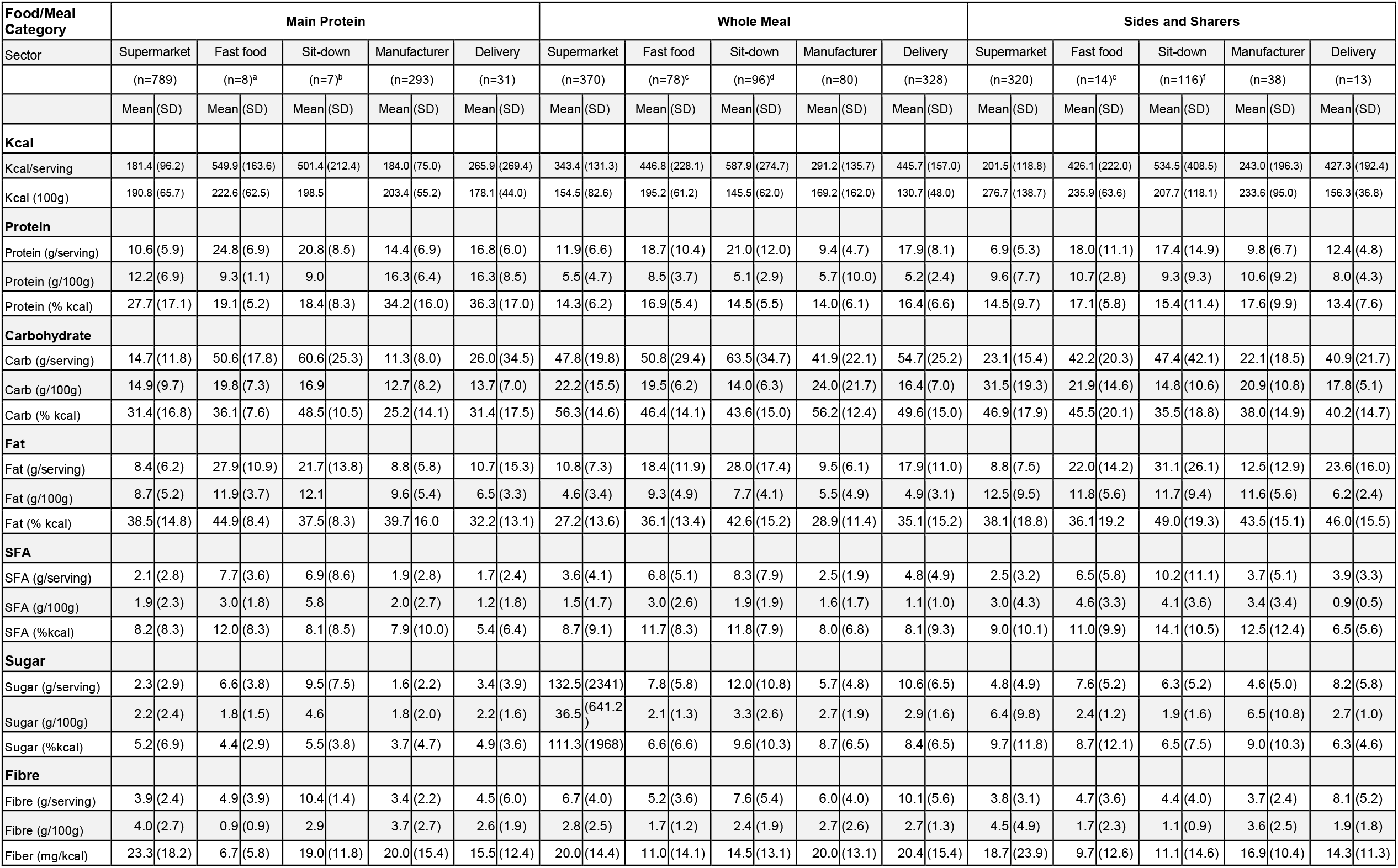
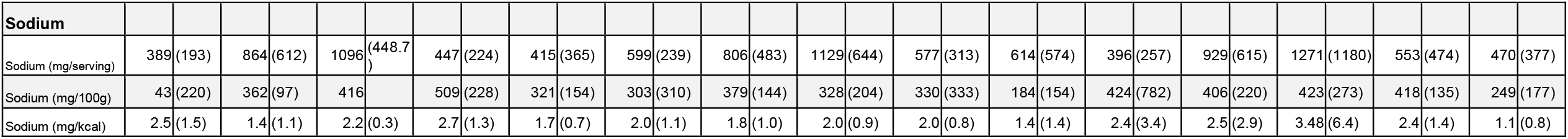
Nutritional composition of the main protein, whole meal and sides and sharers across the US, UK and Canada, combined. ^a^ Nutrition per 100g only available for 3 products; ^b^ Nutrition per 100g only available for 6 products; ^c^ Nutrition per 100g only available for 44 products; ^d^ Nutrition per 100g only available for 71 products; ^e^ Nutrition per 100g only available for 8 products; ^f^ Nutrition per 100g only available for 74 products.

**Table 3:** Nutrition content of meat, vegetarian and vegan items across the US, UK and Canada, combined. ^a^ significantly different from vegan, p<0.001 ^b^ significantly different from vegetarian, p<0.001 ^c^ significantly different from vegetarian, p<0.01

|  | **Meat (n=1507)** | **Vegetarian (n=191)** | **Vegan (n=81)** | **Significance** |
| --- | --- | --- | --- | --- |
| Energy (kcal) | 606 (430-910)<sup>a, b</sup> | 499.0 (370.0-680.0) | 503.0 (345.5-705.0) | <0.001 |
| Protein (g/day) | 35.4 (24.0-51.4)<sup>a, b</sup> | 19.0 (13.0-26.1) | 16.2 (10.5-23.2) | <0.001 |
| Carb (g/day) | 47 (28.7-68.0)<sup>a, b</sup> | 54.0 (38.2-75.6) | 59.0 (38.5-83.5) | <0.001 |
| Sugar (g/day) | 7.0 (4.0-11.0)<sup>a</sup> | 6.9 (3.5-12.9)<sup>a</sup> | 12 (6.0-16.8)<sup>b</sup> | <0.001 |
| Fat (g/day) | 29.0 (18.0-49.0)<sup>a, b</sup> | 23.5 (14.0-33.1) | 19.0 (8.9-31.6) | <0.001 |
| SFA (g/day) | 9.0 (4.6-16.0)<sup>a, c</sup> | 7.0 (4.5-12.0)<sup>a</sup> | 4.0 (2.6-7.4)<sup>b</sup> | <0.001 |
| Fibre (g/day) | 4.0 (2.0-6.0)<sup>a, b</sup> | 5.0 (4.0-7.0)<sup>a</sup> | 8.0 (6.4-11.1)<sup>b</sup> | <0.001 |
| Sodium (g/day) | 1280 (820.0-1952.0) | 865.0 (603.0-1260.0) | 800 (545.0-1410.0) | <0.001 |

#### Protein

Data on protein content was available for 3488 products. The mean protein content per serving (as defined by restaurant or manufacturer) was 11.9 +/- 6.6g (14% kcal) for supermarket main meals; 18.7 ± 10.4g (17% kcal) for fast food main meals; 21.0 +/- 12.0g (15% kcal) for sit down main meals; 9.4 ± 4.7 (14% kcal) for meals available from the manufacturer and 17.9 ± 8.1g (16% kcal) for meals from delivery companies (Table 2). 526 main meals (out of 952) had less than 15g protein per meal. **Figure 2** demonstrates the spread of protein (% kcal) across meal categories.

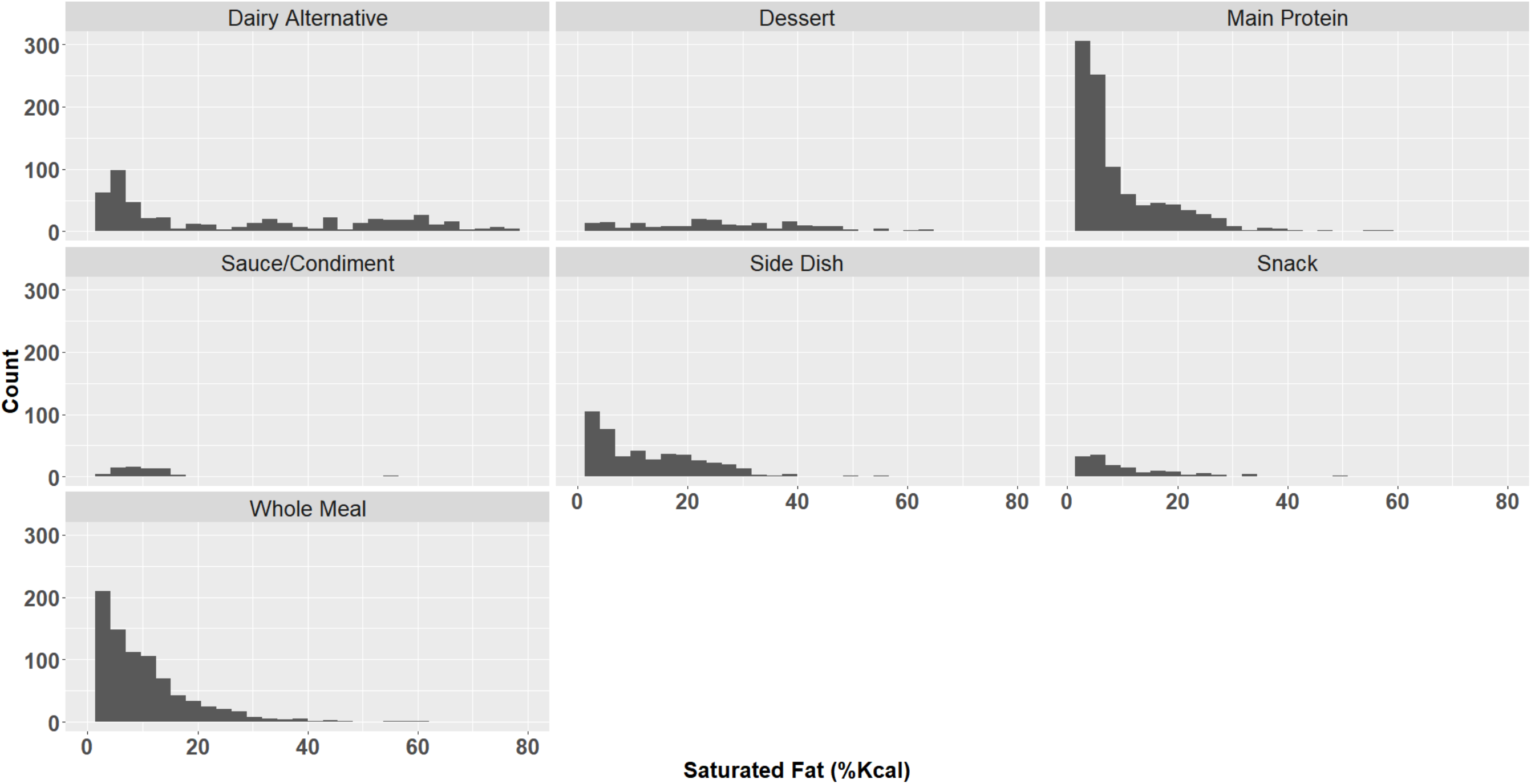

The main protein component per serving was 10.6g ± 5.9 (28% kcal) in supermarkets, 24.8g ± 6.9 (19% kcal) in fast food; 20.8g (18% kcal) in sit-down restaurants; 14.4g ±6.9 (34% kcal) from manufacturers and 16.8g ± 6.0 (36% kcal) from delivery companies. 774 main protein options (out of 1128) had less than 15g per serving.

Sides and sharers provided on average 6.9 ± 5.3g (15% kcal) in supermarkets; 18.0 ± 11.1g (17% kcal) in fast food; 17.4 ± 14.9 (15% kcal) in sit down restaurants; 9.8 ± 6.7g (18% kcal) in manufacturers and 12.4 ± 4.8g (13% kcal) from delivery companies.

#### Carbohydrate

The mean carbohydrate content of main meals varied between 41.9g ± 22.1 (56%kcal) (manufacturers) to 63.5g ± 34.7 (44%kcal) (sit-down restaurants). The mean carbohydrate content of main protein products contained between 11.3g ± 8.0 (25%kcal) (manufacturers) and 60.6g ± 25.3 (49%kcal) (sit-down restaurants) per serving. The fat content of main meals was lowest for the manufacturer sector (9.5g ± 6.1 (29%kcal)) and highest in the sit-down restaurants (28.0g ± 17.4 (43%kcal)).

#### Saturated Fat, Fiber and Sodium

##### Saturated Fat

Saturated fat information was available for 2821 products. The mean %kcal saturated fat of main meals was under 10% kcal for all sectors except for the fast food and sit-down restaurant sectors (11.7% kcal ± 8.3 and 11.8%kcal ± 7.9 respectively). 499 main meals (out of 836) had <10% kcal from saturated fat. Similarly, % kcal saturated fat for the main protein source was under 10% kcal across all sectors except for fast food (12.0% kcal ± 8.3). 678 (out of 973) of the main protein products had <10% kcal from saturated fat. For sides, again, the mean % kcal from saturated was under 10% except for fast food and sit-down restaurants which was 18% kcal and 21% kcal for sides. 214 side products (out of 423) had a % SFA kcal <10%kcal. SFA content demonstrated a striking variability (**Figure 3**) across meal categories, largely due to the use of tropical fats (e.g. coconut milk) driving SFA content of products upward.

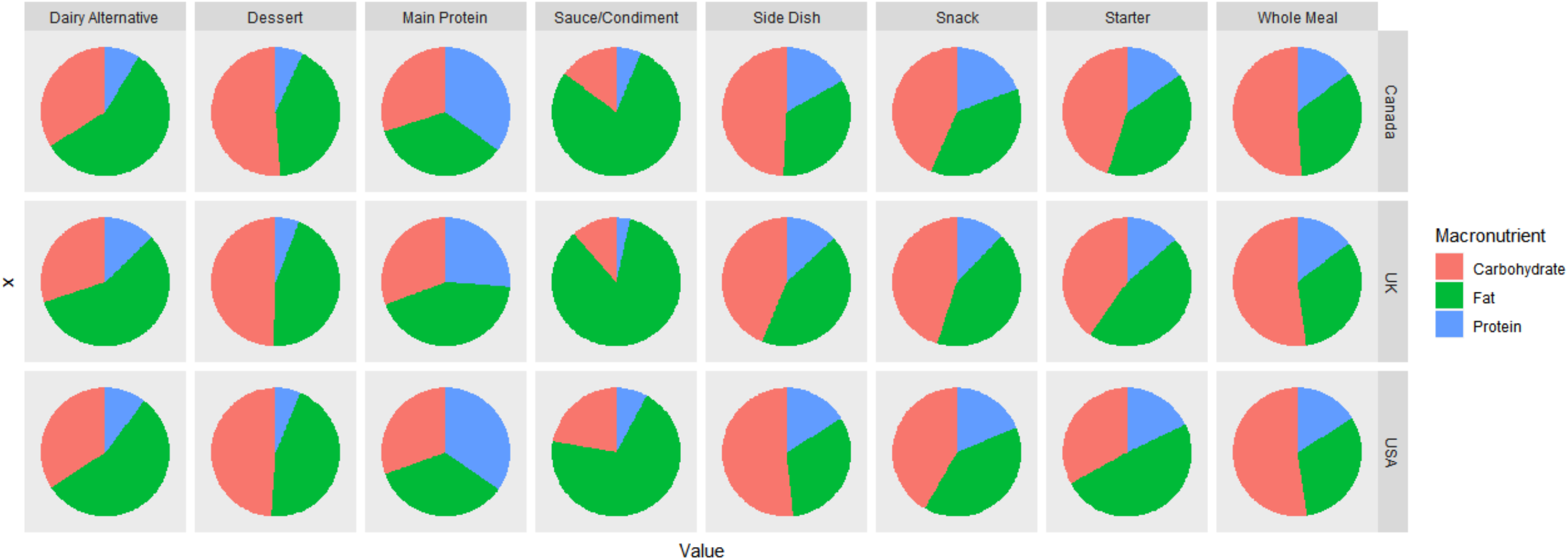

##### Fibre

Data on fibre content was available for 2739 products. The fibre content in main meals ranged from a mean 5.2g ± 3.6 (11mg/kcal ± 14.1) in fast food restaurants to 10.1g +/- 5.6 (20.4mg/kcal ± 15.4) for delivery companies. The main protein had a fibre content ranging from 3.4 ± 2.2 (manufacturers) to 10.4 ± 1.4 (sit-down restaurants). Side/sharers had a fibre content per serving ranging from 3.7 ± 2.4g (manufacturers) to 8.1 ± 5.2g (delivery). 244 main meals (out of 848) had a fibre content of >10g per meal. 25 main protein products (out of 1065) had a fibre content of >10g per meal and 24 side/sharers (out of 456) had >10g per meal.

##### Sodium

Sodium content was available for 3439 items. The sodium content of whole meals ranged from 576 ± 312.9mg (manufacturers) to 1128.8 ± 644.4mg (sit down restaurants) per meal. 790 (out of 922) whole meals had less than 1000mg per meal. The sodium content of the main protein ranged from 389.4 ± 193.1mg (supermarkets) to 1095.7 ± 448.7mg (sit down restaurants). 1001 (out of 1124) main protein products had less than 1000mg sodium. The sodium content for side/sharers ranged from 396.0 ± 256.6mg (supermarkets) to 1270.6 ± 1180.1mg (sit down restaurants). 419 sides/sharers had less than 1000mg (out of 497).

##### Dairy Alternatives

Dairy alternatives represented 700 of the 3488 products. 293 were milks, 166 were cheeses, 83 were yoghurts, with the remaining constituting miscellaneous dressings and dips. The protein content of dairy products was ∼2.5g per serving for both supermarkets and manufacturer sectors. The saturated fat content per serving was 2.5 ± 4.6g (∼20%kcal) for supermarkets and 12.5 ± 7.4 (∼53%kcal) for manufacturers. The fibre content was ∼1g per serving for both sectors, and the sodium content was 125.8 ± 130.1mg for supermarkets and 426.3 ± 233.8mg for manufacturers. There was not sufficient data to assess dairy products from fast food (n=1), sit-down (n=2) and delivery (n=1) sectors.

##### Comparisons to Meat Products

A total of 1657 meat-based products were found across the top ten fast food and sit-down chains in all countries combined. This represented 84% of the total. In contrast there were 219 vegetarian (11%) and 94 (5%) vegan products.

Meat products were significantly higher in energy ((606kcal (IQR: 430-910);(499kcal IQR: 370-680); (503kcal (IQR: 345.5-705kcal)) and protein ((35.4g (24.0-51.4g); 19.0g (13.0-26.1g); 16.2g (10.5-23.2g) compared to the vegetarian (all comparisons = p<0.001) and vegan products (all comparisons = p<0.001), respectively. The meat-based options 9.0g (4.6-16.0g) were also significantly higher in saturated fat than the vegan 4.0g (2.6-7.4g) but not vegetarian 7.0g (4.5-12.0g) options. The meat-based meals were also higher in sodium (1280mg (820-1952mg)) than the vegetarian (865mg (603-1260mg) and vegan (800mg (545-1410mg)).

There were some differences between the vegan and vegetarian options, with the vegan options 4.0g (2.6-7.4) being lower in saturated fat than the vegetarian 7.0g (4.5-12.0g) and higher in fibre (Vegetarian: 5.0g (4.0-7.0g) versus vegan: 8.0g (6.4-11.1g)).

## Discussion

This study aimed to describe and characterise the landscape of plant-based products available across multiple commercial sectors in the US, UK and Canada and document their average nutrition content. We also aimed to compare the number and nutritional content of meat-based, vegan and vegetarian meals available in the largest fast food and sit-down chains across the countries. Overall, the nutritional quality based on pre-defined parameters of the plant-based meals and products was acceptable, and largely met guidelines for protein, saturated fat and fibre content. However, there were some marketed differences between sectors and within both sectors and products which we discuss below.

Plant-based products tend to be marketed as healthy alternatives [23]. In general we found that whole meals available from supermarkets and delivery chains meet dietary requirements for saturated fat content, and the saturated fat content of vegan and vegetarian meals available from both fast-food and sit-down chains was also significantly lower than the meat-based meals. The median saturated fat content of standard UK supermarket main meals has been reported as ∼16%kcal [24], while in the same analysis ready meals specifically marketed as “healthy” have a median saturated fat content of ∼10%kcal [24]. The sodium content of plant-based meals in our study was also low in general, and the mean sodium content per meal was ∼600mg. This is particularly striking as many of the meals were convenience meals such as microwaveable servings which tend to be higher in sodium. Although we did not collect data for meat-based convenience meals in this study for comparison, previous research has found that standard ready meals in the UK contain 890mg (736 to 969mg) per serving [24]. However, the sodium content of plant-based proteins in our analysis was higher than a previous US study examining the sodium content of sentinel foods which found a sodium content of 371mg per serving and 2.3mg per kcal from the 72 items found [25]. The higher concentrations of sodium in our analysis may reflect different inclusion criteria, as we had broader inclusion criteria for plant-based main proteins. Their analysis also included precise measurement of sodium using atomic emission spectroscopy whereas we relied on food labels. Overall, our assessment is therefore that plant-based products - even when convenience-based - might facilitate the consumption of a diet limited in both saturated fat and sodium.

However, we found main meals lower in fibre than we expected. The fibre content of main meals was low for all sectors apart from the specialist delivery sector. At a mean 10g fibre per meal, the delivery meals would provide more than a third of the recommended intake for fibre (28g/day). The main protein component of a meal alone from supermarkets provided about 4g fibre per serving, compared to about 6.5g in a supermarket whole meal. This suggests that the additional ingredients of plant-based supermarket meals consist of fibre-poor starches and limited vegetables, nuts or seeds. This reflects the nature of meat-based supermarket meals which are also low in vegetables and fibre [24, 26].

It is worth mentioning that in general the plant-based whole meals were low to moderate in energy content. Our analysis captured all products marketed as main meals- and this included products marketed as a light lunch, or a full evening meal. The energy content of main meals from supermarkets was only about 350kcal. This therefore offers considerable scope for nutritional intake to be modified by additional foods chosen within the diet.

Dairy alternatives in general offered poor nutritional quality. These products exceeded 10%kcal from saturated fat and were poor sources of protein. This is important to bear in mind for meal and dietary planning. In general, national dietary guidelines recommend two-three servings of dairy per day. For example, the USDA recommends 3 servings of dairy [27], with a serving equalling a cup of milk or yoghurt or 1 to 1.5 ounces of cheese. If dairy foods are exchanged for non-dairy alternatives on a like for like basis, saturated fat intakes based on three servings could reach 15g to 30g from this food group alone depending on the product chosen. However, since most plant-based main proteins and/or plant-based meals are lower in saturated fat than their meat-based alternatives, the relatively high saturated fat content of dairy alternatives may be of no concern. When choosing between dairy products, consumers have the option of easily choosing a high or lower fat version based on clear food labelling. Similarly, making consumers aware that saturated fat may be a particular concern with some plant-based products, especially cheese alternatives, and that some lower fat, or lower saturated fat versions are available (eg, nut based cheeses) will be useful.

The protein content of main meals was around 15%kcal which is within recommended guidelines [8]. However, since again the overall calorie content of meals was low, the absolute content was low, with the majority providing fewer than 15g protein. If rounded up to a 2000kcal diet these products could provide about 65g protein per day - however, since these products are often marketed as a whole meal, it is not clear that the consumer would know to supplement with other protein-rich foods.

In a meat-based diet, meat usually provides the majority of the protein [10]. For this reason, in this analysis we wanted to capture the protein content of the food component being marketed as an alternative to meat-based products. Here, there was considerable variation. The meat protein alternative offered 20 to 25g per serving at fast food and sit-down restaurants, which while lower than their meat-based options, would ensure protein adequacy in most diets. However, the mean protein content of the meal alternative in supermarkets was only about 10g per serving. This average hides some key variation because many products - soy, wheat and pea-protein based provide upwards of 15g per serving, while others, such as jackfruit contain less than 1g. An audit of plant based meat alternatives carried out across supermarkets in Sydney, Australia also found considerable range in the protein content per 100g, for example, from 2.9–20.9g for burgers [15]. These findings have important implications for product labelling and nutritional guidelines which we now discuss.

We deliberately performed our data collection through the lens of a person without expert nutritional knowledge. We wanted to understand which products are visible for a person who is seeking to consume more plant-based products both when in the supermarket, ordering food at home or eating out. From our analysis we believe there are concerns but also opportunities, and these will need to be addressed at the manufacturer, labelling/signposting and nutrition professional levels.

Firstly, as noted above, there are products in supermarkets which are labelled as alternatives to a meat-based main which contain less than 5g of protein per serving. These include products such as jackfruit, vegetable-filled nuggets or patties. In contrast, there are protein-rich products available in supermarkets which would be suitable for a plant-based diet, but are not signposted this way, either via a supermarket website or on the packaging. An example would be a lentil-based pasta, many of which provide >15g protein per serving. Only two legume-based pasta products were identified using our search criteria in one supermarket in the USA. Manufacturers and supermarkets have an important role to play in guiding consumers who are seeking to consume more plant-based products towards products which help them meet their nutritional requirements.

Likewise, dietary recommendations could be clarified to account for the potentially poorer protein content of some plant-based meat alternatives, as suggested by others [14]. For example, the USDA recommends the consumption of 5-7oz (140-200g) of protein equivalents for adult males and females [27]. Currently, it does not address plant-based equivalents made of non-soy protein.

The range of plant-based meat alternatives from pea, wheat and other proteins is increasing markedly and the choice for the consumer has never been greater [2, 3]. Unfortunately, based on our data we have concerns that labelling and dietary guidance is not keeping up. For example, while front-of-pack labelling requires the kcal, fat, saturated fat and salt/sodium content to be specified, it does not require the protein content to be identified. Thus, a person without nutritional knowledge might assume a patty of any kind or any labelled meat alternative would be a nutritional equivalent for their usual meat-based product. We therefore believe there is an opportunity for the manufacturers of protein-rich products (whether plant-based “sausages” or a pasta) to help consumers by specifying the protein content per serving.

Likewise, dietitians and healthcare professionals who provide individual or community based guidance should highlight the issues we have identified above, and give consumers the skills to navigate this rapidly changing landscape. For example, protein requirements could be ensured by including a serving of lentils or other legume with a plant-based meat alternative main, or consuming a soy-based dessert when the main is a vegan pizza or vegan-cheese based pasta.

It is perhaps no surprise that the sector which provided meals and products with the optimal nutritional content in our analysis was the meal-delivery services. By the nature of our search criteria these tended to be companies which were specialist. Some specialised in providing specific types of meals, which included low-carbohydrate, high-protein and vegan, while others were exclusively plant-based meal delivery services. These products were higher in protein and fibre, lower in saturated fat with a moderate sodium content compared to products from the other commercial sectors.

To our knowledge this is the broadest analysis of the plant-based landscape to date. We captured nearly 4000 unique products across three countries, and analysed our data by sector and meal type. Nevertheless, we acknowledge some limitations. Firstly, we relied upon the nutritional information available online for all of the products. As we note in the result section, some products had nutritional information that was identified as being incorrect during the quality control (and was excluded), and we have no definitive way of confirming the accuracy of the information provided. Nevertheless, our findings, both for plant-based and meat-based options are largely in line with other studies including those based on government databases [6, 15, 25].

We used standard prespecified metrics to assess the nutritional quality of the products. The nature and degree of food processing is now recognised as an important factor in the impact of food on health. However, a consensus on grading foods as processed or ultraprocessed is lacking [28, 29], and this information is not readily available on nutritional labels currently. We acknowledge that plant-based products, particularly meat alternatives are processed whereas their meat counterparts are not [30]. However, it’s worth noting that for many plant-based products, their meat-based counterpart would also be processed (eg chicken nuggets, fishcakes etc.).

The aim of our study was to understand the nutritional landscape through the lens of a person without nutritional knowledge, and accordingly only included products which were signposted as being plant-based. For example, if a Chinese restaurant had a tofu-based dish on the menu or a supermarket had a ready meal that was tofu based but neither of them were advertised as being plant-based, then we would not have included these in our analysis. Therefore, we cannot say that the dietary analysis performed here is necessarily a true and accurate representation of all plant-based products available in the countries studied.

We also emphasise that this analysis is a snapshot of the landscape at the time we did the search (April 2020 through December 2020). The plant-based sector is changing rapidly, as noted by the change in the number of plant-based items available in the fast food and sit-down sectors between our two dietary data collection periods. Products are reformulated regularly and therefore the reproducibility of this analysis is limited. Our search was limited to three Western countries, and there is a need for similar studies in other countries and regions.

Finally, we note that the nutritional content of the foods and meals assessed in this study do not necessarily reflect the nutritional content of the whole diet consumed by a person consuming a flexitarian, vegetarian or vegan diet. As others have noted, healthy and unhealthy choices can be made within any dietary pattern [7]. Multiple stakeholders including funders, food scientists, manufacturers, supermarkets, restaurants and public health professionals should work together to ensure food intake from people choosing plant-based products is as healthy as possible.

